# Viromes outperform total metagenomes in revealing the spatiotemporal patterns of agricultural soil viral communities

**DOI:** 10.1101/2020.08.06.237214

**Authors:** Christian Santos-Medellin, Laura A. Zinke, Anneliek M. ter Horst, Danielle L. Gelardi, Sanjai J. Parikh, Joanne B. Emerson

## Abstract

Viruses are abundant yet understudied members of soil environments that influence terrestrial biogeochemical cycles. Here, we characterized the dsDNA viral diversity in biochar-amended agricultural soils at the pre-planting and harvesting stages of a tomato growing season via paired total metagenomes and viromes. Size fractionation prior to DNA extraction reduced sources of non-viral DNA in viromes, enabling the recovery of a vaster richness of viral populations (vOTUs), greater viral taxonomic diversity, broader range of predicted hosts, and better access to the rare virosphere, relative to total metagenomes, which tended to recover only the most persistent and abundant vOTUs. Of 2,961 detected vOTUs, 2,684 were recovered exclusively from viromes, while only three were recovered from total metagenomes alone. Both viral and microbial communities differed significantly over time, suggesting a coupled response to rhizosphere recruitment processes and nitrogen amendments. Viral communities alone were also structured along a spatial gradient. Overall, our results highlight the utility of soil viromics and reveal similarities between viral and microbial community dynamics throughout the tomato growing season yet suggest a partial decoupling of the processes driving their spatial distributions, potentially due to differences in dispersal, decay rates, and/or sensitivities to soil heterogeneity.

## Introduction

Viruses are ubiquitous and abundant members of Earth’s ecosystems that can affect the assembly, dynamics, and function of microbial communities (1–3). They control the size of microbial populations via infection and lysis, redirect microbial metabolism through auxiliary metabolic genes (4–8), and mediate gene transfer across hosts (9,10). In soils, one gram can harbor up to 10^10^ viruses (11,12), sometimes surpassing the number of coexisting bacteria (13). Similar to their role in marine systems (3), recent studies suggest that viruses may be key contributors to carbon and nutrient cycling in terrestrial environments (14–16). Despite this ecological relevance, soil viral communities and the factors shaping them are poorly understood (9,10,17).

Viral replication depends on the successful infection of suitable hosts. As such, the abundances of viral populations and, consequently, the structure of viral communities are inherently linked to the compositional trends of coexisting host communities (18–20). In agricultural soils, rhizosphere processes can alter microbial diversity by actively promoting or inhibiting the recruitment of select taxa (21,22). Soil amendments can further affect microbiome structure by shifting the quantity and quality of available nutrients (23). Understanding whether viral communities display similar trends is essential to unravel the potential of host-virus interactions to affect microbially-influenced soil properties. Additionally, environmental factors such as temperature, pH, nutrient status, and moisture could directly contribute to viral community variation by differentially impacting the activity and decay of soil viruses (24). Similarly, viral adsorption to soil particles could influence the spatial distribution of viruses by limiting virion movement across the soil matrix (24). Thorough characterization of viral diversity patterns across a variety of soil conditions could shed new light on the drivers of viral community dynamics.

Given the absence of a universal marker gene across viral genomes, metagenomic approaches are necessary to survey viral community diversity (25). For soils, recent studies have usually relied on the recovery of viral sequences from whole shotgun metagenomic datasets (14,26,27). This approach not only capitalizes on the existence of streamlined wet lab workflows but also facilitates the simultaneous characterization of the microbial and viral fractions (10). While convenient, most sequences in these total metagenomes are derived from bacterial and eukaryotic genomes (26), presumably concealing the less abundant viral signal unless deep sequencing efforts are performed (10). Moreover, the vast microbial richness associated with soil environments (28) typically results in high-complexity sequence profiles that are challenging to assemble *de novo* (29), further hindering the identification of viral genomes.

Viral communities can be also characterized by physically separating virions from larger microbes through filtration prior to DNA extraction and sequencing. The resulting viral size-fraction metagenomes, or viromes, have increased coverage of viral sequences and can therefore capture a more complete picture of viral diversity relative to total metagenomes (10,30). While this viromic approach has been successfully adopted to study aquatic systems (3,31–33), it has been challenging to implement for soil environments (10), mainly due to low DNA extraction yields that can preclude library construction and sequencing (30). Early soil viromic studies used multiple displacement amplification (MDA) and/or Random Amplified Polymorphic DNA (RAPD) PCR to bypass this limitation (34), however amplification biases, particularly for MDA, preclude quantitative estimations of viral community composition using these approaches (35). Until recently, large amounts of soil input were required to avoid these amplification biases, but recent improvements in library construction from nanograms of DNA now make it practical to work with manageable amounts of soil (∼50 g or less) per sample (30). Continuous development of optimized extraction workflows has enabled virome preparation from a variety of soils (15,36), greatly expanding our ability to examine viral diversity in these environments.

Here, we use a combination of total metagenomes and viromes to characterize soil dsDNA viral communities associated with an agricultural tomato field. First, we compare the performance of the two profiling approaches, in terms of their ability to recover diverse viral sequences, and then we explore viral and microbial ecological patterns in our data. We find that viromes vastly outperform total metagenomes in the recovery of viral diversity and that viral communities display strong spatiotemporal dynamics, which are only partially explainable by shared patterns with bacterial host communities and environmental conditions.

## Materials and Methods

### Sample collection

Samples were collected from a tomato agricultural field in Davis, CA, USA (38°32’08” N, 121°46’22” W) within the context of a larger ongoing study of the impacts of biochar on agricultural production. The soil is classified as Yolo silt loam, a fine-silty, mixed, superactive, nonacid, thermic Mollic Xerofluvents (37). The field was divided into 6.1 m by 4.6 m plots, each with three 1.5 m wide beds. We sampled 8 plots treated with 1 of 4 biochar treatments (650°C pyrolyzed coconut shell [Cool Planet, Greenwood Village, CO, USA], 650°C pyrolyzed pine feedstock [Cool Planet], 800°C pyrolyzed almond shell [Premier Mushrooms and Community Power Corporation, Colusa, CA, USA], and no biochar, **STable 1**) and 1 of 2 nitrogen fertilization regimes (150 or 225 lbs N/acre) (**SFig 1A**). For all sampled plots, 4.214 kg of biochar were buried in the center of each bed on November 8th, 2017; tomato seedlings (cultivar H-8504) were transplanted on May 2nd, 2018; and nitrogen fertilizer was fed through subsurface drip irrigation on five occasions starting May 31st, 2018 (**SFig 1B**). Soil samples from the same 8 plots were collected on April 23rd and August 28th, 2018 for a total of 16 samples.

**Table 1.** Permutational multivariate analyses of variance performed on Bray-Curtis dissimilarities calculated for vOTU profiles in viromes and 16S rRNA gene OTU profiles in total metagenomes (Total MG). The effects of collection time point, biochar, and plot position (coded as column and row within the field) were tested using the whole dataset, while the effect of two different nitrogen amendment amounts was only tested in the August subset of samples, after the amendments had been added.

|  | vOTUs (Virome) |  | 16S OTUs (Total MG) |  |
| --- | --- | --- | --- | --- |
|  | R2 | P | R2 | P |
| Time Point | **0.5** | **0.001** | **0.28** | **0.001** |
| Biochar | **0.139** | **0.042** | 0.166 | 0.331 |
| Nitrogen<br>Amendment<br>(High or Low) | 0.119 | 0.465 | 0.129 | 0.674 |
| Column | **0.173** | **0.001** | 0.048 | 0.419 |
| Row | 0.043 | 0.103 | 0.044 | 0.559 |

**Figure 1.**
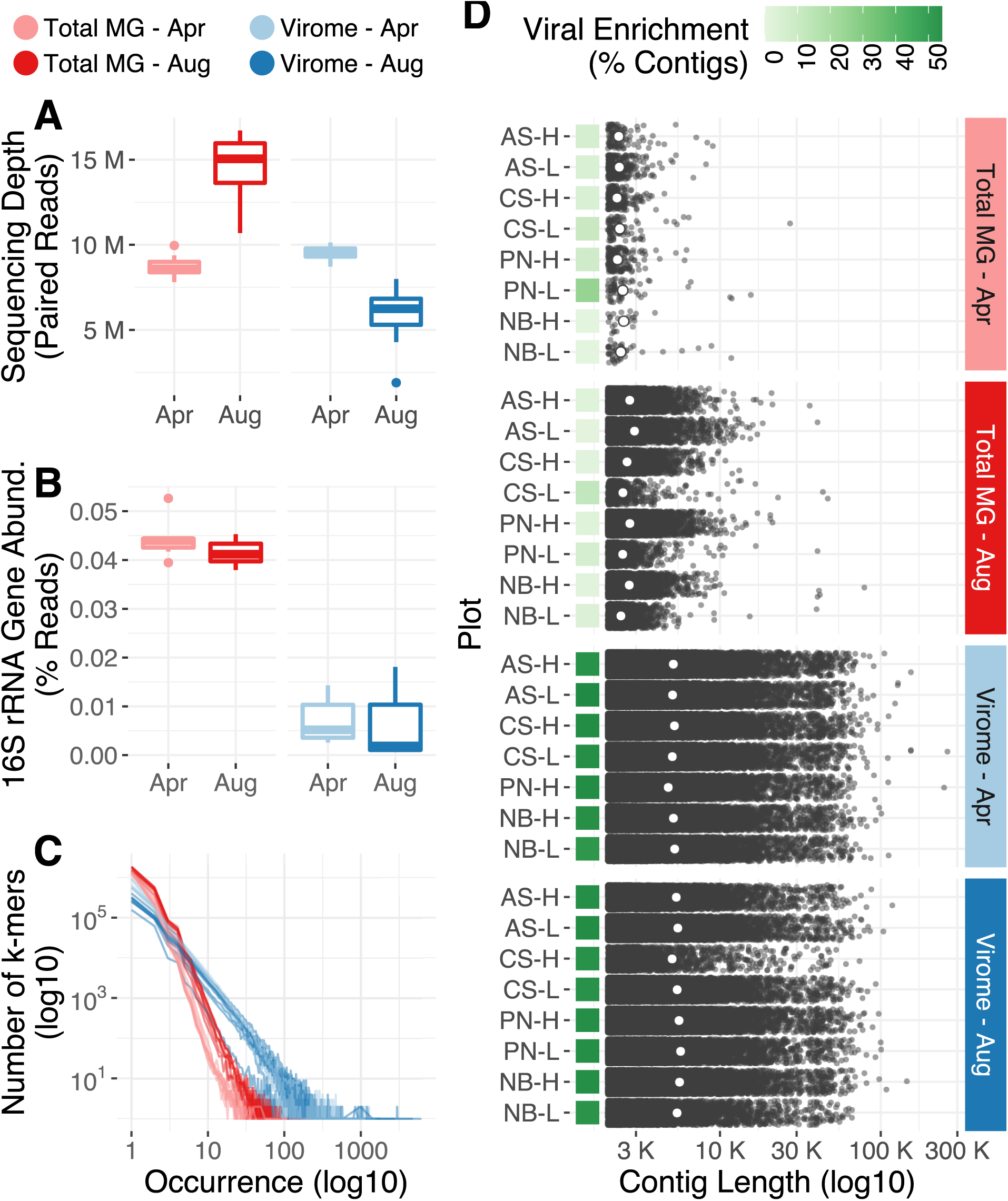
(A) Sequencing depth distribution across profiling methods and collection time points. The y-axis displays the number of paired reads in each library after quality trimming and adapter removal. Boxes display the median and interquartile range (IQR), and data points further than 1.5 x IQR from box hinges are plotted as outliers. (B) Percent of reads classified as 16S rRNA gene fragments in the set of quality trimmed reads; the distribution of data within boxes, whiskers, and outliers is as in panel A. (C) Sequence complexity as measured by the frequency distribution of a representative set of k-mers (k = 31) detected in each library. The x-axis displays occurrence, *i*.*e*., the number of times a particular k-mer was found in a library, while the y-axis shows the number of k-mers that exhibited a specific occurrence. (D) Length distribution of contigs assembled from each library (min. length = 2 Kbp). White dots represent the N50 of each assembly, and green squares display the viral enrichment, as measured by the percent of contigs classified as putatively viral by DeepVirFinder and/or VirSorter.

Soils were harvested near the center of each plot using 2.5 cm soil probes to collect the 0-30 cm depth range. Tomato root systems were present throughout each plot, so August samples were rhizosphere-influenced. A total of 8 soil probes were pooled for each sample, stored in sterile plastic bags, and transported to the laboratory on ice. Within 48 hours, soil samples were sieved to 8 mm and divided for chemistry, moisture, viromics, and total metagenomics.

### Soil chemistry and moisture

Gravimetric moisture content was measured as in Ref (38). Total nitrogen and total carbon were measured using the combustion method, whereas nitrate and ammonium were measured using the flow injection analyzer method in the University of California Davis Analytical Lab (Davis, CA, USA). Soil pH, organic matter, phosphorus (weak Bray and sodium bicarbonate-P), and extractable cations (potassium, magnesium, calcium, and sodium) were measured in the A&L Western Labs (Modesto, CA, USA).

### DNA extractions

A detailed description is provided in the supplementary material. Briefly, for viromics, viral size-fractionation was achieved by resuspending 50 g of soil in AKC’ buffer (39), followed by filtration through a 0.22 μm membrane. Ultracentrifugation was used to concentrate purified virus-like particles, and DNase treatment was used to remove extracellular DNA prior to DNA extraction with the PowerSoil kit (Qiagen, Hilden, Germany). Total DNA for metagenomics was extracted from 0.5 g of soil with the PowerSoil kit.

### Library construction and DNA sequencing

Library construction and high-throughput sequencing were performed by the DNA Technologies and Expression Analysis Cores at the UC Davis Genome Center. Libraries for April samples were prepared with the DNA Hyper Prep library kit (Kapa Biosystems-Roche, Basel, Switzerland) and libraries for August samples were prepared with the Nextera DNA Flex Library kit (Illumina, San Diego, CA) (**SFig 1C**). We were not aware of the differences in library construction methods between sample sets until they became obvious during our bioinformatic processing of the data. Paired-end sequencing (150 bp) was performed across two lanes of the Illumina HiSeq 4000 platform (Illumina), one for each collection time point: viromes and total metagenomes were pooled at equimolar ratios for April samples, and at a 1:2 ratio for August samples. All raw sequences have been deposited in the Sequence Read Archive under the Bioproject accession PRJNA646773.

### Read processing and data analysis

A detailed description is provided in the supplementary material. Briefly, quality-filtering was performed with Trimmommatic (40) and BBDUK (41), followed by *de novo* assembly with MEGAHIT (42) and clustering with PSI-CD-HIT (43). VirSorter (44) and DeepVirFinder (45) were used to detect viral contigs, and vConTACT2 (46) was used to assign taxonomic classifications. Read mapping was performed with BBMap (41), and vOTU coverage tables were generated with BamM (47). Thresholds for defining (>= 10 Kbp, >= 95% global identity) and detecting (>=75% of the contig length covered >=1x by reads recruited at >= 90% average nucleotide identity) viral populations (vOTUs) were implemented in accordance with benchmarking and community consensus recommendations (48,49). Detection and classification of 16S rRNA gene fragments were performed with SortMeRNA (50) and the RDP classifier (51). K-mer profiling was performed with sourmash (52,53). All statistical analyses were done in R using the vegan (54) and DESeq2 (55) packages. All scripts and intermediate files are available at github.com/cmsantosm/SpatioTemporalViromes/

## Results and Discussion

To characterize dsDNA soil viral communities associated with agricultural fields and to better understand the factors that structure them, we collected soil samples from eight tomato plots in Davis, CA, USA (**SFig 1A**). Briefly, each plot was treated with one of four biochar amendments in November 2017: almond shell (AS), coconut shell (CS), pine needle (PN), or no biochar (NB). Tomato seedlings were transplanted in May 2018, and received one of two nitrogen fertilization regimes: 150 lbs N/acre (low, L) or 225 lbs N/acre (high, H). Samples were collected in April (at the pre-planting stage and before any nitrogen additions) and again from the same locations in August to capture two representative stages of one growing season (**SFig 1B**).

To profile the viral communities in these 16 samples and to compare two common techniques for viral community analyses, paired total metagenomes and viral size-fraction metagenomes (viromes) were generated from each sample (with the exception of one April sample, PN-L, that did not yield enough DNA to perform virome sequencing). For both profiling methods, library preparation differed between the two collection time points: April libraries were constructed using a ligation-based approach, while August libraries were constructed with a tagmentation protocol (**SFig 1C**).

### Viromes outperform total metagenomes in the recovery of viral sequences from complex soil communities

To assess the extent to which viral sequences were enriched and bacterial and archaeal sequences were depleted in viromes, relative to total metagenomes, we performed a series of analyses to compare these two approaches. After quality filtering, total metagenomes yielded an average of 8,741,015 paired reads per library for April samples and 14,551,631 paired reads for August samples, while viromes yielded an average of 9,519,518 and 5,770,419 paired reads in April and August, respectively (**Fig 1A, STable 2**). Viromes displayed a significant depletion of bacterial and archaeal sequences, as evidenced by fewer reads classified as 16S rRNA gene fragments: 0.006% of virome reads, compared to 0.042% of reads in total metagenomes (**Fig 1B**). Moreover, taxonomic classification of the recovered 16S rRNA gene reads revealed clear differences in the microbial profiles associated with each approach: total metagenomes were significantly enriched in Acidobacteria, Actinobacteria, Firmicutes, and Thaumarchaeota, whereas viromes were significantly enriched in Armatimonadetes, Saccharibacteria, and Parcubacteria (**SFig 2A-B**). These last two taxa belong to the Candidate Phyla Radiation (CPR) clade and are typified by small cells (56–58), which would be more likely to pass through the 0.22 um filter that we used for viral particle purification (36).

**Figure 2.**
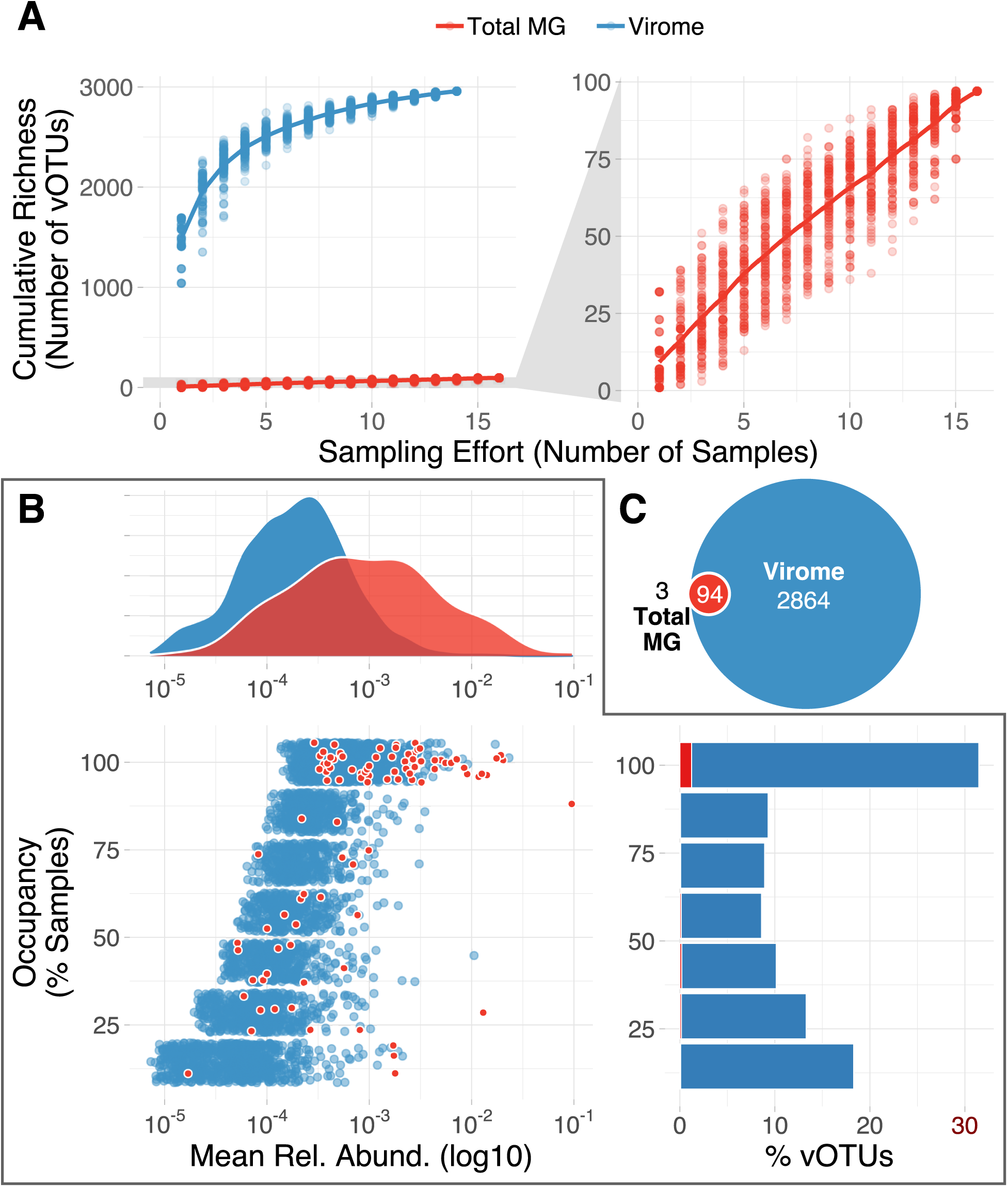
(A) Accumulation curves of vOTUs in total metagenomes (red, n = 16) and viromes (blue, n = 14). Dots represent cumulative richness at each sampling effort across 100 permutations of sample order; the overlaid line displays the mean cumulative richness. The right graph includes the same total metagenomic data as the left graph, zoomed in along the y-axis. (B) Abundance-occupancy data based on vOTU profiles derived from viromes. Data in blue are from vOTUs detected only in viromes, and data in red are from vOTUs detected in both viromes and total metagenomes. Bottom left: Dots represent the mean relative abundance (x-axis) and occupancy (percent of samples in which a given vOTU was detected, y-axis) that individual vOTUs displayed in viromes within a collection time point (April or August). Thus, vOTUs detected in both time points are represented twice. Red dots highlight the set of vOTUs shared between total metagenomes and viromes. Top: Density curves showing the distribution of relative abundances for all vOTUs detected in viromes (blue) or the subset of vOTUs detected in viromes and total metagenomes (red). Bottom right: Percent of vOTUs (x-axis) found at each occupancy level (y-axis). Red boxes highlight the percent of vOTUs detected in both profiling methods. (C) Euler diagram displaying the overlap in detection for each vOTU (n = 2,961) across profiling methods. Red vOTUs were detected by both profiling methods, and three vOTUs were detected exclusively in total metagenomes.

To assess differences in sequence complexity between the two profiling methods, we calculated the k-mer frequency spectrum for each library (**Fig 1C)**. Relative to viromes, total metagenomes displayed an increased number of singletons (k-mers observed only once) and an overall tendency towards lower k-mer occurrences, indicating that size-fractionating our soil communities reduced sequence complexity. These differences in sequence complexity translated into notable contrasts in the quality of *de novo* assemblies obtained from individual libraries (**Fig 1D**): while viromes yielded 800 Mbp of assembled sequences across 169,421 contigs (250 Mbp assembled in ≥ 10 Kbp contigs), total metagenomes produced only 65 Mbp across 22,951 contigs (1.5 Mbp assembled in ≥ 10 Kbp contigs). The improved assembly quality from the viromes was despite lower sequencing throughput relative to total metagenomes, particularly for the August samples (**Fig 1A**). Using DeepVirFinder (45) and VirSorter (44) to mine assemblies for viral contigs, we found that 52.4% of virome contigs and only 2.2% of total metagenome contigs were identified as viral. Together, these results show that our laboratory methods for removing contamination from cells and free DNA reduced genomic signatures from cellular organisms, substantially improved sequence assembly, and successfully enriched the viral signal in soil viromes relative to total metagenomes.

### Viromes facilitate exploration of the rare virosphere

To remove redundancy in our assemblies, we clustered all 192,372 contigs into a set of 105,909 representative contigs (global identity threshold = 0.95). Following current standards to define viral populations (48,49), we then screened all non-redundant ≥10 Kbp contigs for viral signatures. We identified 4,065 viral operational taxonomic units (vOTUs) with a median sequence length of 17,870 bp (max = 259,025 bp) and a median gene content of 27 predicted ORFs (max = 421 ORFs). To profile the viral communities in our samples, we mapped reads against this database of non-redundant vOTU sequences (≥ 90% average nucleotide identity, ≥ 75% coverage over the length of the contig). On average, 0.04 % of total metagenomic reads and 23.4% of viromic reads were mapped to vOTUs (**SFig 3A**). One August virome sample (CS-H) had particularly low sequencing throughput and low vOTU recovery (**Fig 1A** and **SFig 3B**) and was discarded from downstream analyses.

**Figure 3.**
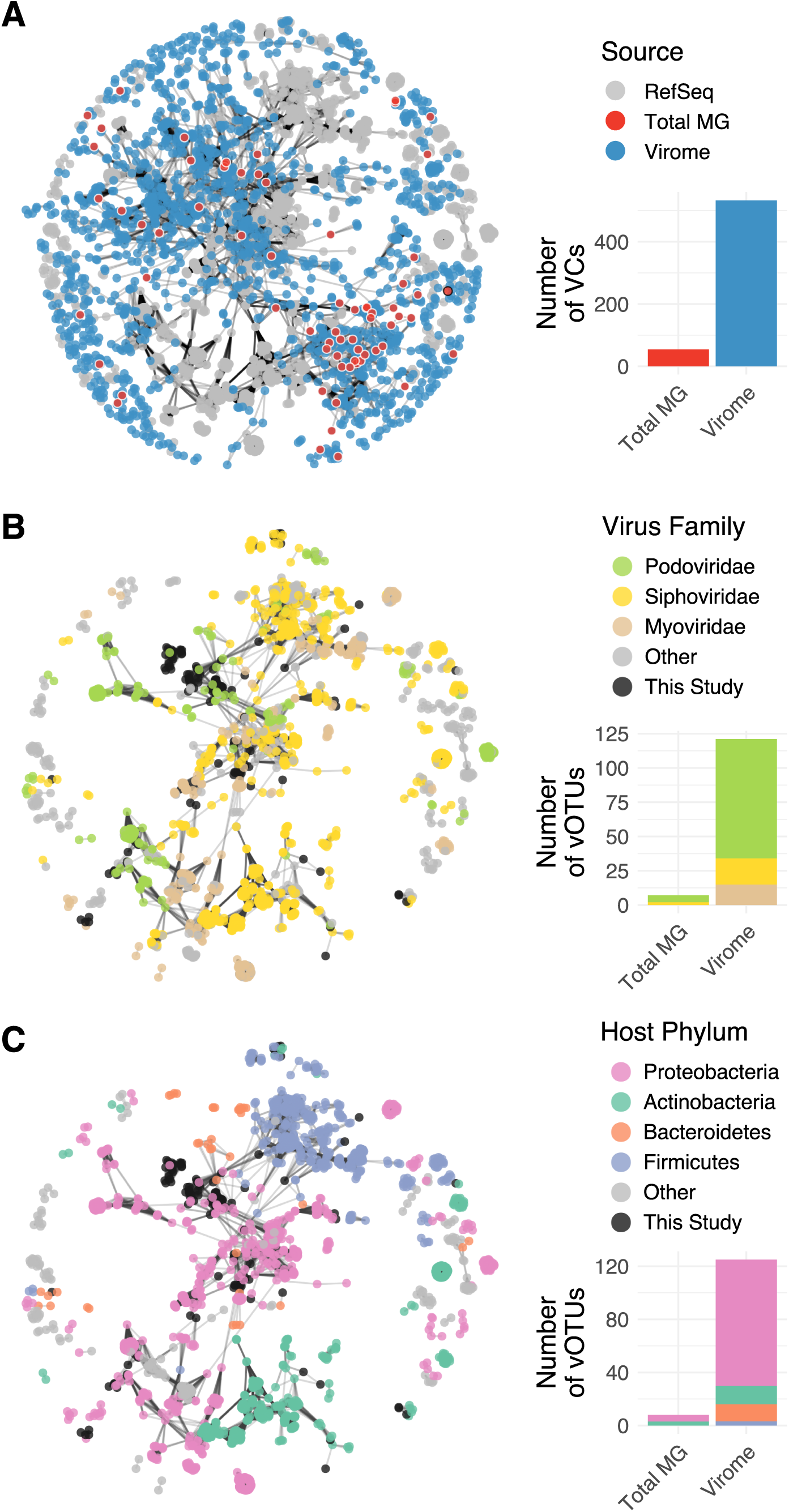
(A) Gene-sharing network of vOTUs detected in viromes alone (blue nodes), total metagenomes (red nodes; nodes outlined in white were also detected in viromes, nodes outlined in black were detected exclusively in total metagenomes), and RefSeq prokaryotic virus genomes (grey nodes). Edges connect contigs or genomes with a significant overlap in predicted protein content. Only vOTUs and genomes assigned to a viral cluster (VC) are shown. Accompanying bar plots indicate the number of distinct VCs detected in total metagenomes and viromes (VCs detected in both profiling methods are counted twice, once per bar plot). (B, C) Subnetwork of all RefSeq genomes and co-clustered vOTUs. Colored nodes indicate the virus family (B) or the associated host phylum (C) of each RefSeq genome. Barplots display the number of vOTUS classified as each predicted family (B) or host phylum (C) across total metagenomes and viromes (vOTUs detected in both profiling methods are counted twice, once per bar plot).

In total, 2,961 vOTUs were detected through read mapping in at least one sample. Of these, 2,864 were exclusively found in viromes, 94 in both viromes and total metagenomes, and 3 in total metagenomes alone (**Fig 2C**). Thus, viromes were able to recover thirty times as many viral populations as total metagenomes, even when vOTUs assembled from viromes were part of the reference set for read mapping. Consistent with capturing a representative amount of viral diversity from the viromes but not total metagenomes, our sampling effort was sufficient to generate a saturation plateau in vOTU accumulation curves derived from viromes but not total metagenomes (**Fig 2A**).

To examine the distribution of vOTUs along the abundance-occupancy spectrum, we used the viromic data to compare mean relative abundances of vOTUs with the number of virome samples in which each vOTU was detected. Given the contrasting experimental conditions between the April and August samples, we performed this analysis within each time point. Highly abundant vOTUs tended to be recovered in the majority of virome samples (*i*.*e*., they displayed high occupancies), while rare vOTUs were typically recovered in only a few, a trend typically observed in microbial communities (59). Moreover, abundance-occupancy patterns for the 94 vOTUs detected in both viromes and total metagenomes revealed that vOTUs recovered from total metagenomes were among the most abundant and ubiquitous in the virome profiles (**Fig 2B**), suggesting that total soil metagenomes are biased towards the recovery of highly prevalent vOTUs and are more likely to miss the rare virosphere.

### Viromes reveal a diverse taxonomic landscape

To examine the taxonomic landscape covered by our vOTUs, we compared them against the RefSeq prokaryotic virus database using vConTACT2, a network-based method to classify viral contigs (46). Under this approach, vOTUs are grouped by shared predicted protein content into taxonomically informative viral clusters (VCs) that approximate viral genera. Of the 2,961 vOTUs, 1,712 were confidently assigned to VCs, while the rest were only weakly connected to other clusters (outliers, 784 vOTUs) or shared no predicted protein content with any other contigs (singletons, 465 vOTUs) (**STable 3**). Only 130 vOTUs were grouped with RefSeq genomes, indicating that this dataset has substantially expanded known viral taxonomic diversity (**Fig 3A**). Subsetting the vOTUs detected by each profiling method showed that viromes captured a more taxonomically diverse set of viruses: 1,711 vOTUs detected in viromes were assigned to 533 VCs, while 68 vOTUs detected in total metagenomes (67 of which were also detected in viromes) were assigned to 54 VCs (53 of which were also detected in viromes). This suggests that concerns about biases in the types of viruses recovered through soil viromics (*e*.*g*., through preferential recovery of certain viral groups from the soil matrix) may be unfounded and/or that any such biases also apply to total metagenomes, at least for the viruses and soils examined here.

Most of the 130 vOTUs clustered with RefSeq viral genomes could be taxonomically classified at the family level (**Fig 3C**). Podoviridae was the most highly represented family, followed by Siphoviridae and Myoviridae. Myoviridae were only detected in viromes, not total metagenomes (**Fig 3B**), further confirming that viromes do not seem to exclude viral groups relative to total metagenomes–if anything, the opposite seems to be true. Among the Siphoviridae clusters, we could further identify three vOTUs as belonging to the genus Decurrovirus, which are phages of *Arthrobacter*, a genus of Actinobacteria common in soil (60–62). Because the genome network was highly structured by host taxonomy (**Fig 3C**, (63)), we used consistent host signatures among RefSeq viruses in the same VC to assign putative hosts to vOTUs in VCs shared with RefSeq genomes. Most such vOTUs were putatively assigned to Proteobacteria, Actinobacteria, or Bacteroidetes hosts, and a few were linked to Firmicutes. Interestingly, these bacterial phyla were among the most abundant taxa in the 16S rRNA gene profiles derived from the total metagenomes from these soils (**SFig 2A**).

Although soil viromes and total metagenomes have been qualitatively compared and their presumed advantages and disadvantages have been reviewed (10), this is the first comprehensive comparison of results from both profiling approaches applied to the same samples. Here we show that soil viromes clearly recover richer (**Fig2A**) and more taxonomically diverse (**Fig3**) soil viral communities than total metagenomes.

### Compositional patterns of agricultural soil viral communities and their ecological drivers

Since viromes vastly outperformed total metagenomes in capturing the viral diversity in our samples, we focused on viromes to explore compositional relationships among viral communities. We first performed a permutational multivariate analysis of variance (PERMANOVA) on Bray-Curtis dissimilarities to assess the impacts of collection time point, spatial location, biochar treatment, and nitrogen amendments on beta diversity (**Table 1)**. Differences between April and August viral communities were the most significant, explaining 50.0% of the variance in our dataset. Interestingly, the position of each sampled plot along the west-east axis, *i*.*e*. the field column in which each plot was located (**SFig 1A**), was the second most important factor (17.3% of total variance) in separating viral communities. Biochar treatment accounted for 13.9% of the variance, but its effect was only borderline significant (p = 0.048). Finally, since nitrogen additions occurred after the April samples had been collected, we tested the effect of the two different fertilizer concentrations (150 or 225 lbs N/acre) on the composition of viral communities in August only, and found no significant effect (**Table 1**).

To assess whether the bacterial and archaeal communities displayed similar compositional trends and could therefore potentially explain patterns in viral community composition, we attempted to generate metagenome assembled genomes (MAGs) from our total metagenomes. However, the low quality of total metagenomic assemblies (**Fig1D**) precluded MAG reconstruction (19 MAGs with a median completeness of 30.3), so instead we used 16S rRNA gene profiles recovered from total metagenomes (**SFig 2A**). Although 16S rRNA genes accounted for less than 0.05% of the reads in our total metagenomes (**SFig 1B**), 573 OTUs were detected, and richness asymptotes were attained in accumulation curves (**SFig 2C**), suggesting that enough microbial diversity was recovered to justify further analyses. A PERMANOVA on Bray-Curtis dissimilarities revealed that only collection time point significantly correlated with bacterial and archaeal community composition, while spatial location, biochar treatment, and nitrogen fertilizer concentration did not (**Table 1**). Below, we further examine these temporal and spatial patterns, along with their relationships with measured soil chemical properties.

### Viral and microbial communities display coupled temporal dynamics

Compositional differences between April and August samples were significant for both viral and microbial communities (**Fig 4A**), and a Mantel test revealed that the Bray-Curtis dissimilarity matrices of viral and microbial communities were significantly correlated (R = 0.59, P = 0.0003). Unsurprisingly, due to the reliance of viruses on their hosts for replication, viral and microbial communities have been previously observed to correlate in soil (14,18), and results here suggest that viruses and their bacterial and archaeal hosts had coupled temporal responses to the same variables and/or to each other.

**Figure 4.**
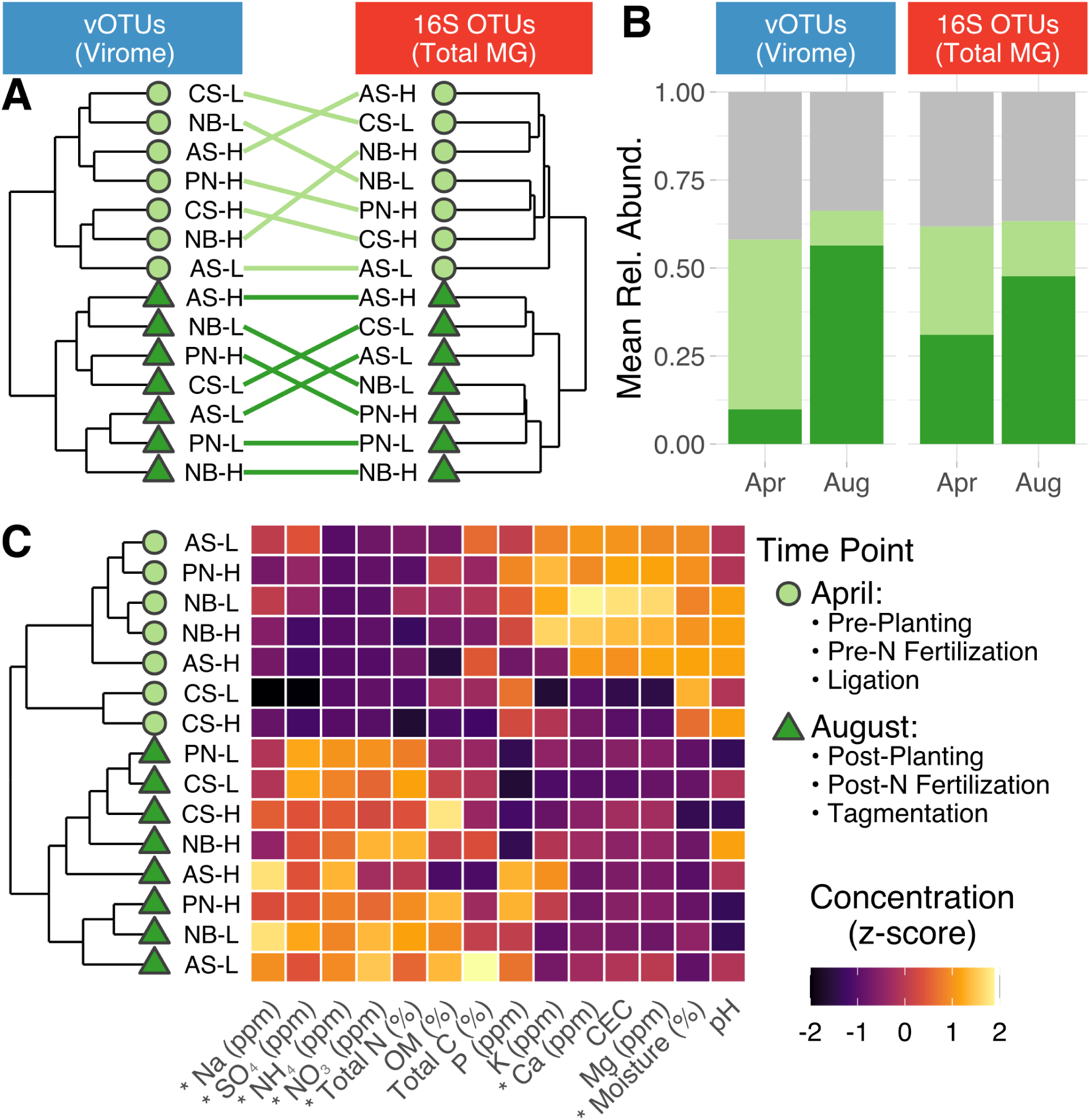
(A) Tanglegram depicting the hierarchical clustering of viral communities (left) and bacterial and archaeal (hereafter, microbial) communities (right) based on Bray-Curtis dissimilarities. Shapes and colors indicate collection time point (legend to the right of panel C). Colored lines connect viral and microbial communities from the same sample. (B) Summed mean relative abundances of the set of vOTUs and 16S rRNA gene OTUs significantly affected by collection time point. Color indicates enrichment in April (light green), August (dark green), or no significant enrichment by time point (grey). (C) Hierarchical clustering of samples based on soil chemical profiles and environmental characteristics. The heatmap shows the z-transformation of each measurement across samples. OM = organic matter, CEC = cation exchange capacity.

To identify individual vOTUs that displayed temporal dynamics, we performed a differential abundance analysis between collection time points. Of 2,958 vOTUs detected in viromes, 1,062 were enriched in April samples, and 681 were enriched in August samples (**STable 4**). The summed relative abundances of these vOTUs accounted for up to 63.3% of the viral communities (**Fig 4B**). No clear taxonomic trend could be detected, as very few (92 of 1,743) differentially abundant vOTUs could be taxonomically classified, and all identified viral families and putative host phyla were proportionally represented across April- and August-enriched vOTUs (**SFig 4A-B**).

To identify microbial taxa associated with compositional shifts between time points, we performed a similar differential abundance analysis on the 16S rRNA gene profiles. Of 573 total OTUs, 38 were enriched in April, and 20 were enriched in August (**STable 5**). This relatively small number of OTUs accounted for 66.3% of the total microbial community abundance (**Fig 4B**). OTUs enriched in April encompassed a diverse set of taxa, with Bacteroidetes, Alphaproteobacteria, Gammaproteobacteria, and Acidobacteria being the most represented, while OTUs enriched in August were exclusively members of the Actinobacteria or Alphaproteobacteria (**SFig 4C**). Interestingly, a recent study on greenhouse-grown tomatoes (64) showed similar taxonomic enrichment and depletion patterns in rhizosphere communities relative to bulk soil communities, including enrichment for Actinobacteria and Alphaproteobacteria in the rhizosphere, suggesting that root microbiome recruitment could explain some of the compositional changes between pre-planting and harvest time points. These results are consistent with many previous studies that have demonstrated rhizosphere impacts on the diversity and composition of microbial communities (65–68) and, more recently, on RNA viral communities in Mediterranean grasslands (16).

We also inspected soil properties and nutrient profiles to determine whether they followed the same temporal patterns as viral and microbial communities. Indeed, they did: samples grouped into two distinct clusters based on overall chemical composition, one with all April samples and one with all August samples. (**Fig 4C**). Testing the impact of collection time point on individual chemical measurements revealed that, relative to April soils, August soils exhibited a significant increase in ammonium, nitrate, total nitrogen, sodium, and sulfate concentrations and a significant decrease in soil moisture and calcium content (ANOVA, **STable 6**). Nitrogen amendments, which were applied between sampling time points, have been shown to shift the composition of microbial communities associated with tomato rhizospheres towards an Actinobacteria-enriched state (23), suggesting that the increased abundance of this phylum in August samples may be related to an increase in nitrogen availability, in addition to the presence of tomato roots. Furthermore, a recent survey of rice paddies revealed that nitrogen amendments influenced the absolute abundances of viruses and bacteria (69), indicating that a coupled response by viral and microbial communities could have been triggered by agricultural nitrogen inputs. Considering that soil moisture availability is one of the main factors influencing the composition of soil bacterial communities (70) and that soil desiccation has been linked to viral inactivation (24), it is likely that soil moisture also contributed to viral and bacterial community differences between time points. Similar correlations between viral community composition and soil moisture content have been observed in thawing permafrost peatlands (14).

Altogether, these results indicate that viral and microbial communities display strong temporal dynamics that are consistent with responses to changes in both biotic (plant) and abiotic characteristics of soils. Still, we acknowledge that the magnitude of this temporal shift could have been amplified by biases associated with library preparation: while April samples were constructed using a standard ligation-based approach, August samples were inadvertently constructed with a tagmentation protocol due to changes in methodology at the sequencing facility (see Methods). Given that non-random transposition might skew the compositions of viromic (71) and metagenomic profiles (72), it is possible that some of the detected temporal differences in our study could stem from technical artifacts. However, considering the consistency of some of our results with previous studies (*e*.*g*. specific taxonomic response patterns associated with rhizosphere assembly and nitrogen fertilization), it is likely that, overall, the observed compositional differences represent true ecological patterns over time.

### Viral communities but not microbial communities were spatially structured across an agricultural field

To visualize the effect of plot position on the overall structure of viral communities, we performed a principal coordinate analysis on Bray-Curtis dissimilarities calculated across the virome-derived vOTU profiles. While the first axis captured the differences between collection time points, the second axis revealed a compositional transition from the westernmost (W) to the easternmost (E) end of the field (**Fig 5A-B**). Comparing the pairwise Bray-Curtis dissimilarities in viral community composition against the corresponding pairwise W-E field distances further confirmed a significant correlation between viral community composition and spatial location (**Fig 5C**). We found 1,035 vOTUs displaying a spatial gradient across the field: 460 with increasing abundances from east to west and 575 with increasing abundances from west to east (**Fig 5D, STable 7**). Furthermore, examining the overlap with vOTUs affected by collection time point (**Fig 4B**) revealed that more than half of the spatially structured vOTUs were temporally dynamic (**STable 7, Fig 5D**), suggesting that spatial structuring of viral communities occurred on time scales captured by this study.

**Figure 5.**
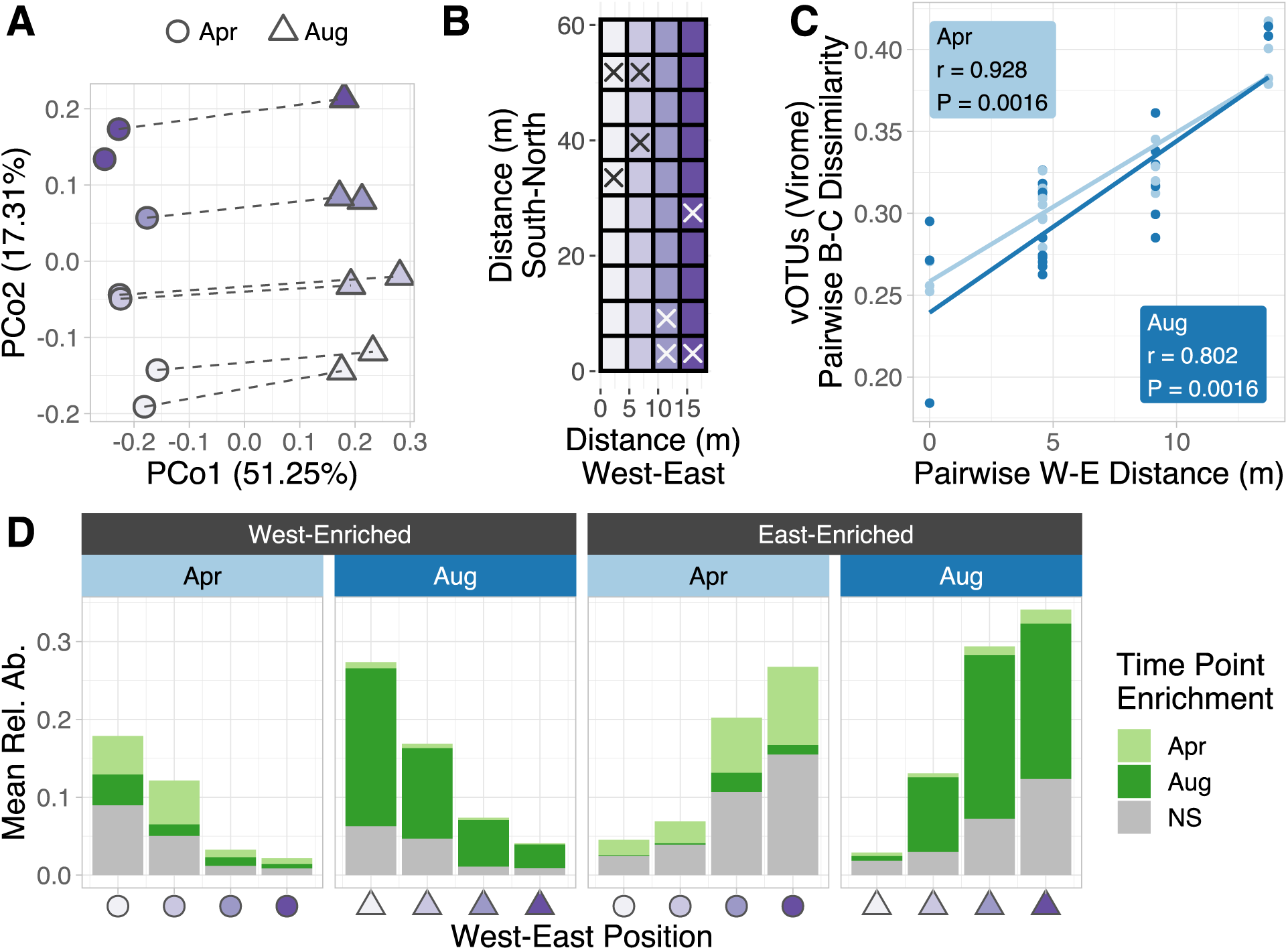
(A) Unconstrained analysis of principal coordinates based on vOTU Bray-Curtis dissimilarities calculated across virome samples. Shapes represent the collection time point, and colors indicate the position of each plot along the west-east axis of the sampled field, as displayed in B. Lines connect April and August samples collected from the same plot. (B) Diagram depicting the spatial distribution of plots. Sampled plots are indicated by an “X” (C) Correlations between spatial distance across the west-east axis (in meters between plots) and Bray-Curtis dissimilarities of viral communities profiled in viromes. Inset values display the r statistic and associated P-value of Mantel tests performed within each collection time point. (D) Shifts in the mean summed relative abundances of significantly West-enriched or East-enriched vOTUs across the W-E axis in April or August. Bar colors indicate whether vOTUs were also significantly enriched in a particular time point (NS = Not Significant).

We next explored whether this spatial gradient was observed in the bacterial and archaeal communities. As expected from the PERMANOVA results (**Table 1**), there was no significant correlation between microbial community composition and spatial distance across the W-E field axis (**SFig 5A-B**). Thus, while overarching and/or temporal differences in viral and microbial communities were related, the spatial distribution of viral and microbial communities would seem to result from at least partially decoupled assembly processes. This result is consistent with Mantel tests between viral and microbial community composition, which were significant overall (*i*.*e*., driven by temporal separation, see above), but were not significant within each collection time point (April: R = 0.12, P = 0.269; August: R = 0.27, P = 0.149). We also analyzed the soil nutrient profiles and did not detect a significant effect of W-E position on any of the measured chemical or environmental properties (**STable 6**).

Given the strong spatiotemporal dynamics displayed by viral communities, we examined the effect of biochar treatment on overall viral community composition through a partial canonical analysis of principal coordinates (CAP) that removed the variance due to collection time point and W-E position. This approach revealed that biochar-amended soils clustered separately from untreated soils and that viral communities associated with different types of biochar were compositionally distinct, a pattern driven by 43 differentially abundant vOTUs (**SFig 6A**,**C, STable 8**). Similar trends were observed in the 16S rRNA gene profiles of total metagenomes despite the effect of biochar on overall microbial community composition being not significant (**SFig 6B, Table 1**). None of the measured chemical soil properties were significantly influenced by biochar treatment (**STable 6**). Overall, these results suggest that biochar amendments may have impacted both viral and microbial community composition but that temporal differences (and, for viral communities, spatial differences) were far more pronounced. Thus, future studies that seek to investigate the impacts of soil amendments on viral communities may benefit from rigorous spatiotemporal replication in the study design.

While spatial differentiation has been previously reported for soil viral communities, prior studies considered larger areas with contrasting soil properties. A RAPD DNA study of soil viral diversity along a 4 km land-use transect (including forest, pasture, and cropland systems) found significant differences in viral community composition along the transect (73). Similarly, a metagenomic characterization of peatland soils found distinct viral communities in three areas comprising different habitats along a permafrost gradient within a ∼12,500 m^2^ area (14). Although the spatial patterns in viral community composition detected in these studies correlated with differences in various environmental parameters, none of the soil properties that we measured in this study were able to explain the observed spatial gradient. Moreover, we found no evidence of spatial structuring in the coexisting bacterial and archaeal communities, suggesting that viruses were uniquely affected by an unknown underlying factor.

One possible explanation for the observed spatial structuring of viral communities could be unidentified legacy effects from previous growing seasons that may have created an unrecognized gradient across the agricultural field. It has been proposed that rates of viral decay might be much slower than viral production in some soils (74), such that an accumulation of older viruses might explain differences in the spatial structuring of viral, relative to microbial communities. If this were the case, we would expect most of the spatially structured vOTUs to persist over time and exhibit the same abundance patterns in the April and August samples, yet more than half of these vOTUs were also differentially abundant between time points. Another potential factor that could have contributed to this spatial differentiation is the location of the sampled field next to a gravel road. Although traffic along this road is minimal (it is inside a fenced, university-owned agricultural research field site), it is conceivable that vehicular traffic could differentially disperse viruses across the field. Additionally, contrasting conditions at the border of the field could also have led to edge effects. While little is known about the impact of edge effects on soil viruses, other biotic and abiotic properties of agricultural soils have been shown to be influenced by their proximity to the field border (75,76). Future studies could be designed to better disentangle the complex interactions of production, decay, selection, and dispersal on viral and microbial community composition.

## Conclusions

By comparing total metagenomes and viromes from the same tomato soil samples, we showed that viromes recovered a richer and more taxonomically diverse set of vOTUs than total metagenomes, even at lower sequencing depths. Moreover, total metagenomes mostly detected only the highly persistent and abundant vOTUs, a pattern that highlights the greater ability of viromes to explore the rare virosphere. Analyzing the beta-diversity trends of viral and microbial communities revealed coupled temporal shifts that coincided with changes in the biotic and abiotic properties of soil. Viral communities further displayed a compositional gradient along the sampled agricultural field that was not observed in the microbial communities, suggesting that there are factors that differentially affect the spatial distribution of viral and microbial communities. These strong spatiotemporal patterns eclipsed a minor effect of biochar treatment on viral community structure, emphasizing the importance of replicated experimental design over space and time when evaluating the effect of treatments on soil viral communities in future experiments.

## Acknowledgements

This work was supported by the UC Davis College of Agricultural and Environmental Sciences and Department of Plant Pathology (new lab start-up to JBE). Support for CSM was provided by an award from the U.S. Department of Energy, Office of Science, Office of Biological and Environmental Research, Genomic Science Program, Number DE-SC0020163. Funding for the field experimental setup and agricultural activities was provided by the California Department of Food and Agriculture Fertilizer Research and Education Program, Award #16-0662-SA and the Almond Board of California, Award #17-ParikhS-COC-01-0 (grants to SJP). The sequencing was carried out at the DNA Technologies and Expression Analysis Core at the UC Davis Genome Center, supported by NIH Shared Instrumentation Grant 1S10OD010786-01

