## Supplemental Methods and Figures for "Viromes outperform total metagenomes in revealing the spatiotemporal patterns of agricultural soil viral communities"

### Supplementary Material

### Supplementary Methods

*Soil viral DNA extraction*

Viral DNA was extracted from 50 g of fresh soil per sample using a protocol modified from Ref. (1). For each sample, two 50 ml conical tubes were filled with 25 g of soil and 37.5 ml of 0.02 µm filtered AKC' extraction buffer (per liter: 10% PBS, 10g K Citrate, 1.44 g $Na_2HPO_4$, 0.24g $KH_2PO_4$, and 36.97g $MgSO_4$). Resulting slurries were vortexed briefly until homogenized, shaken on an orbital shaker for 15 minutes at 400 RPM, vortexed for 3 minutes, and centrifuged at 4,700 x g for 15 minutes. Supernatant was filtered through a 0.02 µm filter to remove most cells, and supernatants from tubes corresponding to the same sample were combined into one 70 ml polycarbonate ultracentrifuge tube. Supernatants were centrifuged using an Optima LE-80K ultracentrifuge (Beckman-Coulter Life Sciences, Indianapolis, IN, USA) with a 45 Ti rotor at 32,000 x g for 3 hours at 4 °C.

Supernatant was decanted and the resulting pellets (the viral fraction), were resuspended in 200 µl of ultrapure water. Extracellular DNA was removed from the resuspended pellets by treatment with 30 units of RQ1 RNase-free DNase and 30 µl of 10X DNase buffer (Promega Corp., Madison, WI, USA) for two hours at room temperature (22 °C). The reaction was quenched with 30 µl of DNase stop solution (Promega Corp., Madison, WI, USA).

DNA was extracted from the viral fraction using the DNeasy PowerSoil kit (Qiagen, Hilden, Germany), following the manufacturer's protocol, except the bead-beating step was replaced by a 10-minute 70 °C incubation, a 5-second vortex, and a 5-minute 70 °C incubation. Extracted DNA was quantified by an Invitrogen Qubit 4 Fluorometer using a 1x High Sensitivity DNA assay (Thermo Fisher Scientific, Inc., Waltham, MA, USA).

*Soil total DNA extraction*

Total DNA from soil was extracted from 0.5 g of soil using the DNeasy PowerSoil kit (Qiagen, Hilden, Germany), following the manufacturer's instructions. Extracted DNA was quantified by an Invitrogen Qubit 4 Fluorometer using a 1x High Sensitivity DNA assay (Thermo Fisher Scientific, Inc., Waltham, MA, USA).

*Read processing and assembly*

Trimmomatic (2) was used to remove library adapters and quality-trim raw reads (minimum q-score of 30 evaluated on 4-base sliding windows; minimum read length of 50). BBDuk (3) was then used to remove PhiX sequences. *De novo* assembly on individual libraries was performed with MEGAHIT (4) in meta-large mode with a contig cutoff size of 2,000 bp. Assembly statistics were generated using the BBTools stats.sh script (3). To remove redundant contigs across assemblies, we used the PSI-CD-HIT (5) implementation of BLASTN to cluster contigs at a global identity threshold of 0.95.

*Detection and classification of viral contigs*

We used VirSorter (6) and DeepVirFinder (7) to identify putative viral contigs. For VirSorter, only contigs assigned to categories with the most confident (categories 1 and 4) or likely (categories 2 and 5) predictions were retained; for DeepVirFinder, only sequences with a score ≥ 0.9 and p-value < 0.05 were retained. The union of contigs identified by both methods was used for downstream analyses. Viral contig identification was first performed on individual assemblies to measure the enrichment of viral signal in the set of contigs found in each library. Viral contig identification was also performed on the subset of clustered contigs with lengths ≥ 10 Kbp to generate a database of non-redundant sequences representing viral operational taxonomic units (vOTUs). The length threshold and 95% sequence identity from prior clustering were based on previous benchmarking and definitions of vOTUs (8,9).

To assign taxonomic classifications to the recovered vOTUs, we first predicted protein content using Prodigal in metagenome mode (10). The generated amino acid file was then used to build a gene-sharing network using vConTACT2 (11) with the following parameters: NCBI RefSeq database of bacterial and archaeal viral genomes (v85) was used as a reference, Diamond (12) was used to calculate protein alignment, and MCL (13) and ClusterOne (14) algorithms were used to calculate protein and genome clusters, respectively.

*Read mapping*

Read mapping against the database of non-redundant vOTUs was performed with BBMap (3) at a minimum sequence identity of 90%. Resulting SAM files were converted to BAM files and indexed using SAMtools (15). The parse function of BamM (16) was then used to generate two vOTU tables: one displaying the trimmed pileup coverage (tpmean mode) and the other one displaying the absolute number of mapped reads (counts mode). Finally, we calculated the per-sample horizontal coverage for each vOTU using the genomecov function in BEDtools (17). We then identified instances in which vOTUs displayed < 75% coverage over the length of the contig and filtered them out of the vOTU tables using an in-house R script (github.com/cmsantosm/SpatioTemporalViromes/blob/master/Processing/Scripts/votu_filtering.Rmd)

*Detection and classification of 16S rRNA gene fragments*

Reads containing 16S rRNA gene sequences were recovered using SortMeRNA (18) by comparing quality-filtered reads against representative versions of the bacterial and archaeal SILVA databases (19). The RDP classifier (20) was then used to assign taxonomy. The output hierarchical file was further parsed using the hier2phyloseq function implemented in the RDPutils package (21) to generate a count table.

*K-mer profiling*

Sourmash (22,23) was used to compute k-mer signatures for each library using a compression ratio of 1,000 and k-mer size of 31.

*Data analysis and visualization*

All statistical analyses were conducted using R version 3.6.3. Unless otherwise stated, analyses were performed using the trimmed pileup coverage vOTU table. The vegan package (24) was used for the following analyses: accumulation curves were calculated using the specaccum function, Bray-Curtis dissimilarity matrices were calculated on Hellinger-transformed relative abundances using the vegdist function, permutational multivariate analyses of variance (PERMANOVA) were performed with the adonis function, and Mantel tests were performed with the mantel function. The function pcoa from the package ape (25) was used to perform principal coordinate analyses. Hierarchical clustering on z-transformed values was performed with the hclust function. Differential abundance analyses were performed with DESeq2 (26) using count tables as input. All plots were generated with the ggplot2 (27), ggdendro (28), GGally (29), and eulerr (30) packages. All scripts and intermediate files are available at github.com/cmsantosm/SpatioTemporalViromes/

### Supplementary Figures

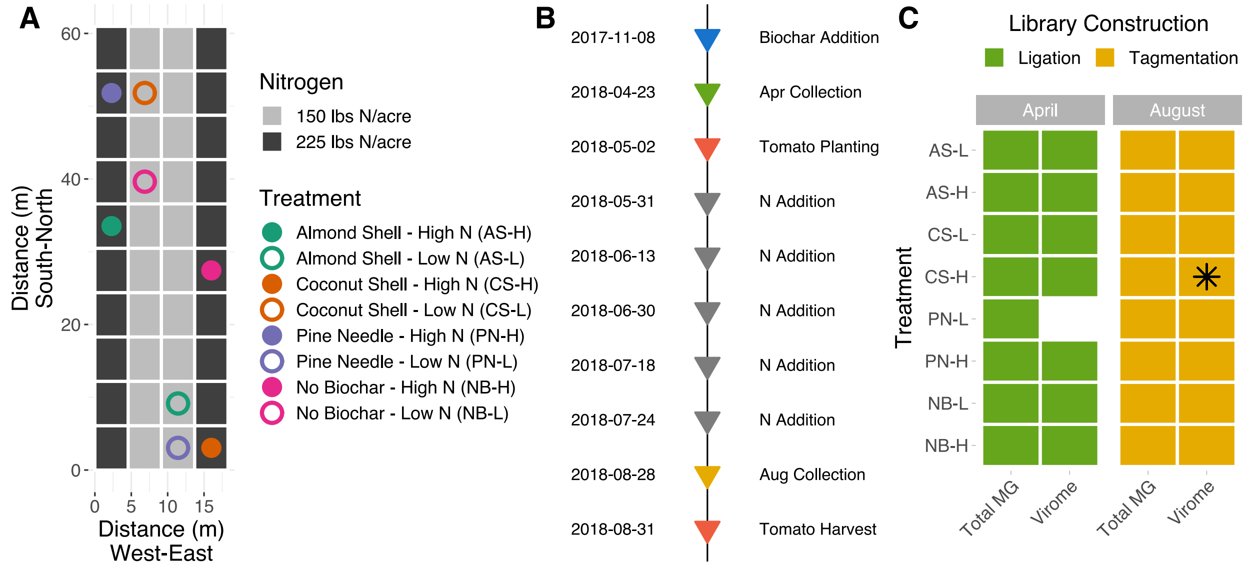


**Supplementary Figure 1**

(A) Diagram depicting the spatial distribution of plots in the agricultural field. Dots indicate the eight sampled plots with color corresponding to the biochar treatment and fill or outline corresponding to the nitrogen fertilization treatment (low - outline only, high - filled). Fertilization began between the two collection time points (see B). (B) Timeline of the growing season, biochar and nitrogen amendments, and collection time points. (C) Library construction workflows used to generate the total metagenomes (MG) and viromes. The blank space represents a virome sample that failed at the library construction step, and the asterisk highlights the virome sample omitted from compositional analyses due to suboptimal sequencing throughput and read mapping (see Figures 1A and D and Supplementary Figure 3).

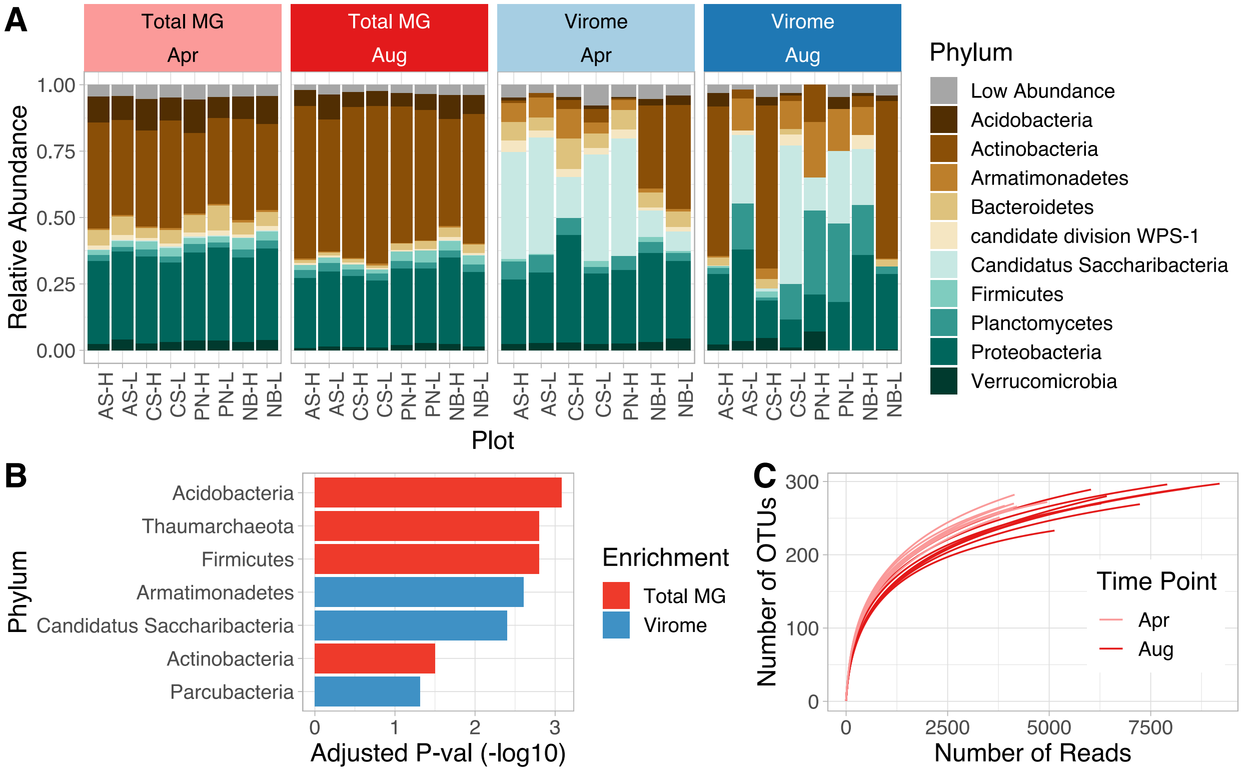


**Supplementary Figure 2**

(A) Relative abundances of microbial phyla in each library, based on recovered 16S rRNA gene fragments with an assigned taxonomy. Only the top 10 most abundant phyla are shown. “Low abundance” represents the summed relative abundances of all OTUs not assigned to the 10 most abundant phyla. (B) Microbial phyla with significantly different relative abundances between total metagenomes and viromes (adjusted $P < 0.05$; paired Wilcoxon test). Red and blue bars indicate phyla significantly enriched in total metagenomes and viromes, respectively. Higher values along the x-axis indicate more significant enrichment. (C) Rarefaction curves of 16S rRNA gene OTU profiles derived from total metagenomes. Each line represents one metagenome.

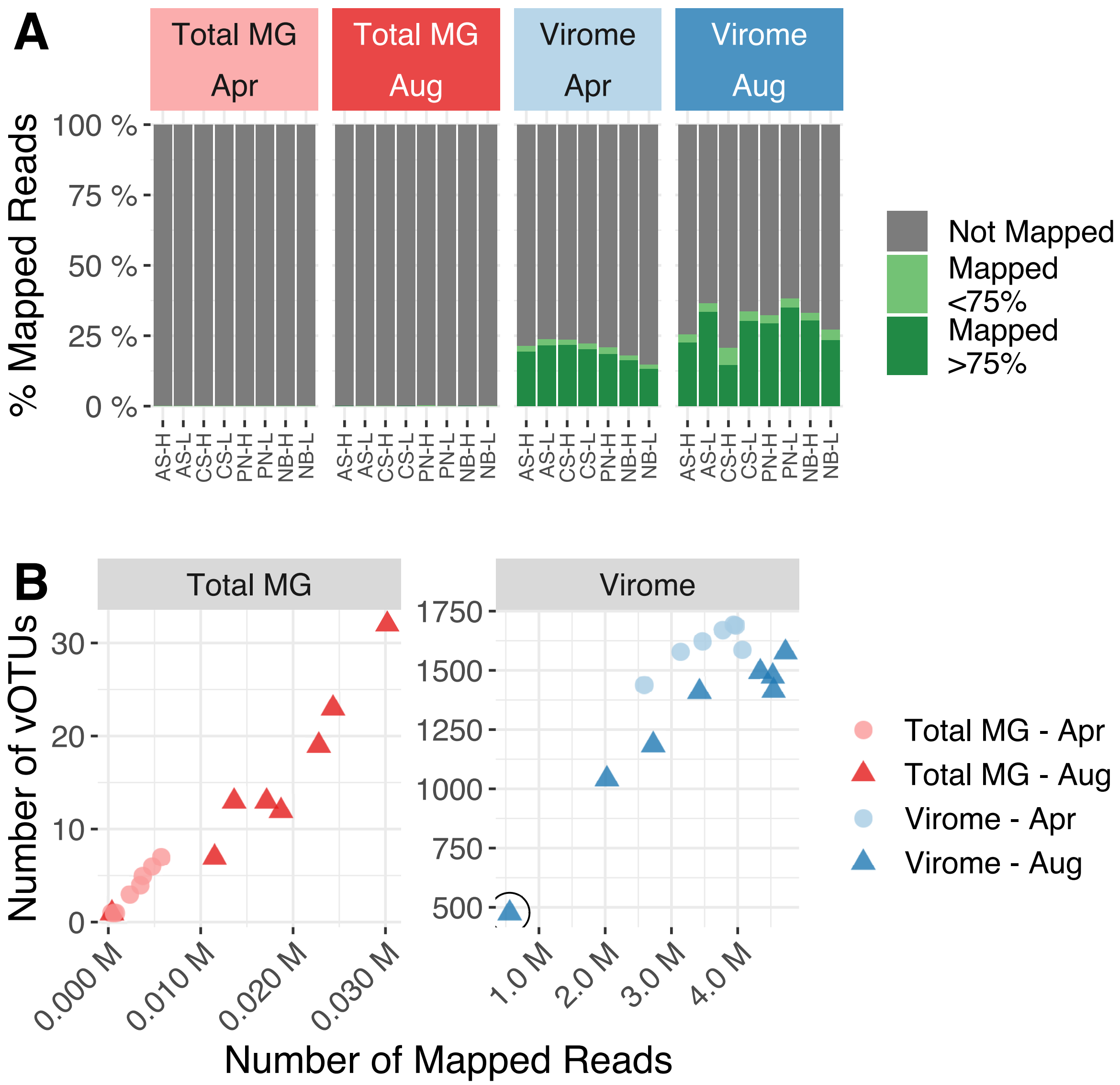


**Supplementary Figure 3**

(A) Percent of high-quality reads mapped to the set of vOTU contigs identified in our dataset. Green colors highlight read mapping below (light green) or above (dark green) the threshold of 75% coverage over the length of the vOTU sequence required for detection. (B) Distribution of the total number of reads mapped to vOTUs (x-axis) and the total number of vOTUs detected in each sample (y-axis). Note different y-axis maxima between graphs. The circled virome sample performed suboptimally (>2 standard deviations below the mean number of mapped reads and mean richness across viromes) and was therefore discarded for downstream compositional analyses.

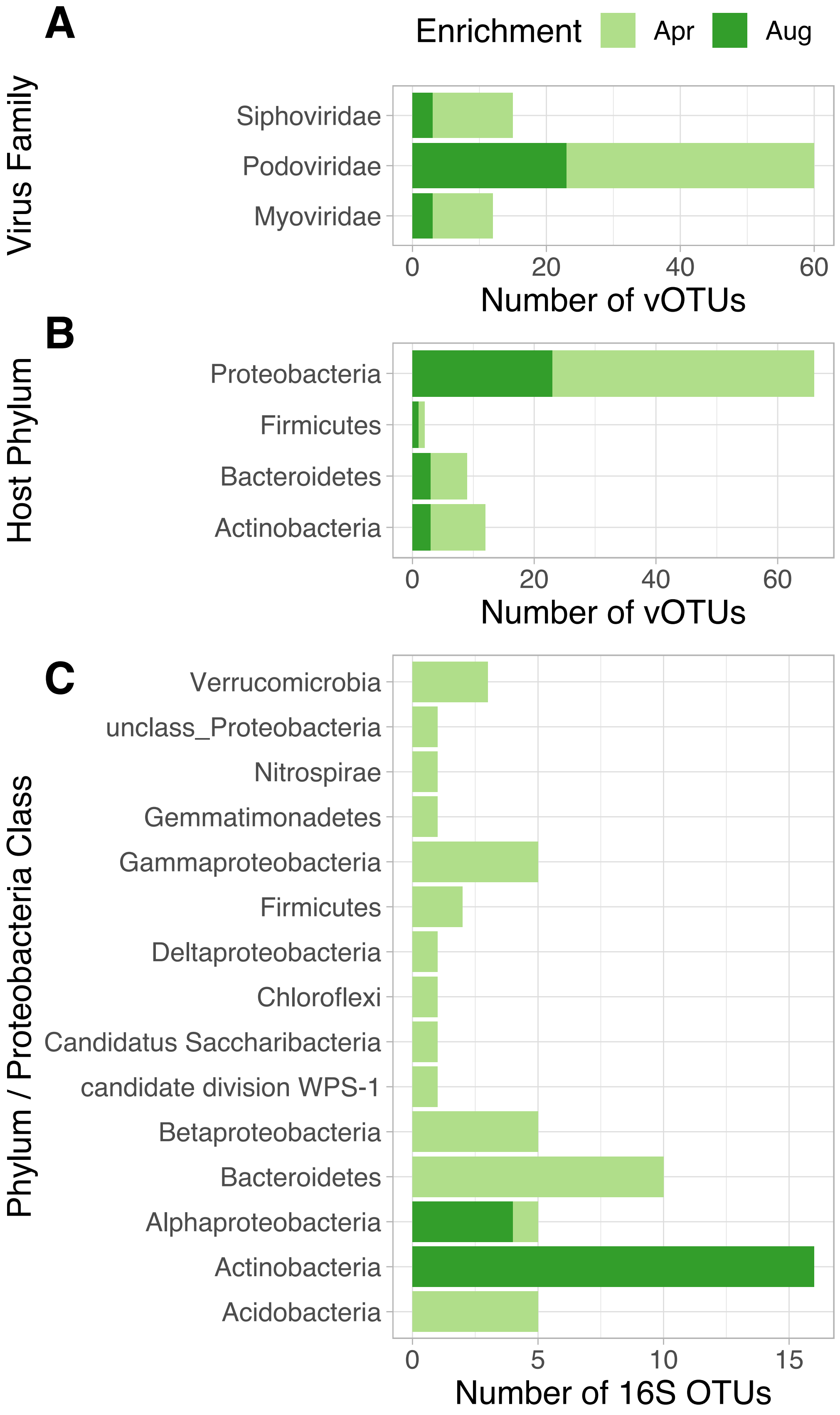
A
B
C
Enrichment
Apr
Aug
Virus Family
Siphoviridae
Podoviridae
Myoviridae
0
20
40
60
Number of vOTUs
Host Phylum
Proteobacteria
Firmicutes
Bacteroidetes
Actinobacteria
0
20
40
60
Number of vOTUs
Phylum / Proteobacteria Class
Verrucomicrobia
unclass_Proteobacteria
Nitrospirae
Gemmatimonadetes
Gammaproteobacteria
Firmicutes
Deltaproteobacteria
Chloroflexi
Candidatus Saccharibacteria
candidate division WPS-1
Betaproteobacteria
Bacteroidetes
Alphaproteobacteria
Actinobacteria
Acidobacteria
0
5
10
15
Number of 16S OTUs

216

**Supplementary Figure 4**

(A) Virus family and (B) predicted host phylum of vOTUs that were significantly enriched according to collection time point and that could be taxonomically classified (n = 92 vOTUs) (**Fig 3**). (C) Phylum or Proteobacteria class of the 16S rRNA gene OTUs that were significantly differentially abundant between collection time points. In all plots, color indicates the collection time point in which vOTUs or 16S rRNA gene OTUs were enriched.

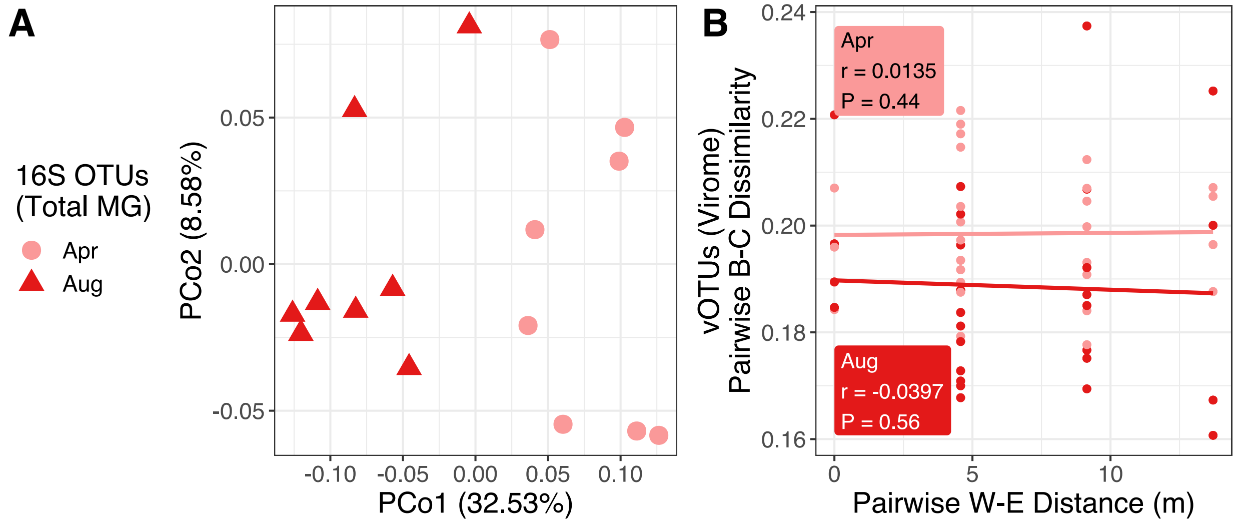


**Supplementary Figure 5.**

(A) Unconstrained analyses of principal coordinates based on Bray-Curtis (B-C) dissimilarities calculated across 16S rRNA gene OTU profiles derived from total metagenomes. (B) Correlation between spatial distance across the west-east axis (in meters between plots) and Bray-Curtis dissimilarities calculated across 16S rRNA gene OTU profiles derived from total metagenomes. Inset values display the Mantel r statistic and associated P-value.

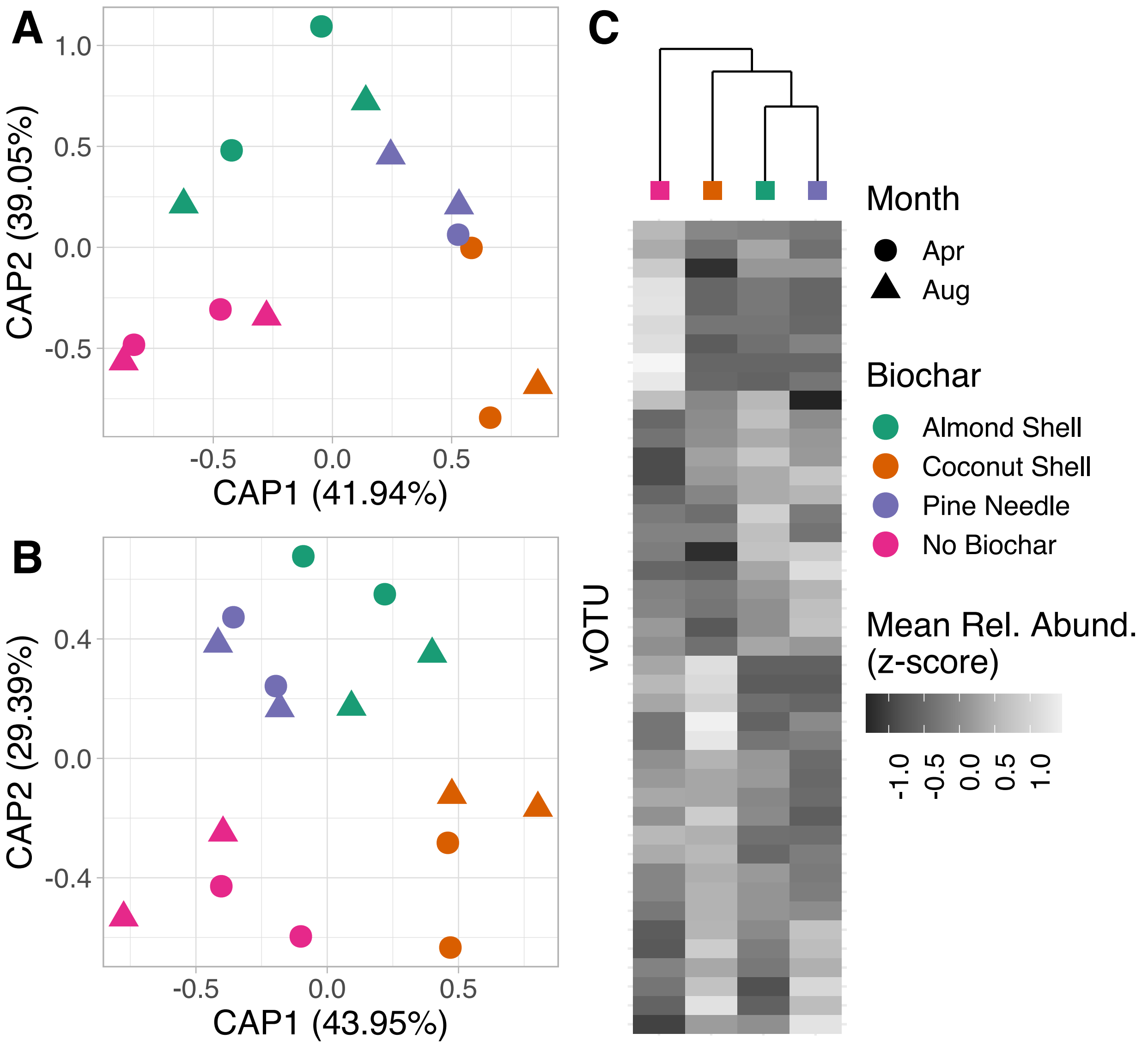


**Supplementary Figure 6.**

(A) Partial canonical analysis of principal coordinates (CAP) performed on Bray-Curtis dissimilarities calculated on vOTU profiles from viromes. The effects of collection time point and W-E position were removed. Colors indicate biochar treatment, and shape indicates sampling time point (legends are to the right of panel C).(B) Partial CAP performed on Bray-Curtis dissimilarities calculated on 16S rRNA gene OTU profiles from total metagenomes. The effect of collection time point was removed. (C) Hierarchical clustering of biochar treatments based on the relative abundances of vOTUs significantly affected by biochar amendments. The heatmap shows the mean relative abundance (z-transformed) of each vOTU (rows) across biochar treatments (columns).

### Supplementary Table Legends

**Supplementary Table 1**

Properties of biochar amended to agricultural field

**Supplementary Table 2**

Sequencing depths (before and after quality filtering) and sample metadata for all libraries reported in this study.

**Supplementary Table 3**

Viral cluster assignment for RefSeq genomes and vOTU contigs

**Supplementary Table 4**

Set of vOTUs differentially abundant across collection time points (Wald test, adjusted P-val < 0.05)

**Supplementary Table 5**

Set of 16S rRNA gene OTUs differentially abundant across collection time points (Wald test, adjusted P-val < 0.05)

**Supplementary Table 6**

ANOVA test of the effects of collection time point, biochar, nitrogen amendment concentration, and plot position (coded as column and row within the field) on the measured chemical properties of soil. The effect of nitrogen amendment concentration was only tested for the August samples.

**Supplementary Table 7**

Set of vOTUs significantly affected by plot position along the west-east axis of the sampled field  (Wald test, adjusted P-val < 0.05)

**Supplementary Table 8**

Set of vOTUs differentially abundant across biochar treatments (likelihood ratio test, adjusted P-val < 0.05)
